# Root system growth and function response to soil temperature in maize (*Zea mays* L.)

**DOI:** 10.1101/2023.03.15.532822

**Authors:** Randy Clark, Dan Chamberlain, Christine Diepenbrock, Mark Cooper, Carlos D. Messina

## Abstract

Crop adaptation to the mixture of environments that defines the target population of environments is the result from a balanced resource allocation between roots, shoots and reproductive organs. Root growth places a critical role in the determination of this balance. Root growth and function responses to temperature can determine the strength of roots as sinks but also influence the crop’s ability to uptake water and nutrients. Surprisingly, this behavior has not been studied in maize since the middle of the last century, and the genetic determinants are unknown. Low temperatures often recorded in deep soil layers limit root growth and soil exploration and may constitute a bottleneck towards increasing drought tolerance, nitrogen recovery, sequestration of carbon and productivity in maize. High throughput phenotyping (HTP) systems were developed to investigate these responses and to examine genetic variability therein across diverse maize germplasm. Here we show that there is: 1) genetic variation of root growth under low temperature and below 10°C, and 2) genotypic variation in water transport under low temperature. Using simulation, we demonstrate that the measured variation for both traits contribute to drought tolerance and explain important components of yield variation in the US corn-belt. The trait set examined herein and HTP platform developed for its characterization reveal a unique opportunity to remove a major bottleneck for crop improvement, and adaptation to climate change.

## Introduction

When root system water supply does not meet leaf transpiration demand, water deficits and stress occur, and a plethora of molecular pathways, hormonal signals, and physiological responses are activated (Reynolds et al., 2021; Karlova et al., 2021). These morphological and hydraulic coordinated responses are not fully understood (Maurel and Nacry, 2020). In maize (*Zea mays* L.), a symptom known as leaf rolling becomes visible within hours of the onset of water stress due to decreasing water potential and turgor within the leaves (Baret et al., 2018). An example of this phenomenon was observed in 2013 in breeding research trials in Elgin, Nebraska where measurable available soil water was still being recorded (Fig. 1), eliciting questions of why a symptom of water deficit was observed in the presence of available soil water; whether low water leaf potentials could be underpinned by limited root exploration and/or water transport; how low temperature affects root occupancy and water transport in maize; and to what degree does genetic or genotypic variation exists for these traits.

**Figure 1.**
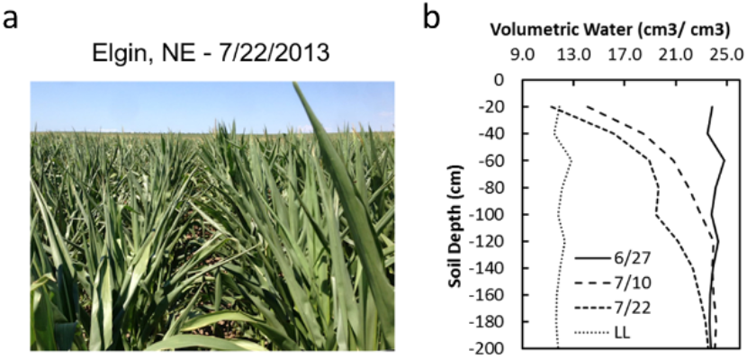
Expression of leaf rolling in field in a maize crop (a) with remaining available soil water as estimated by volumetric soil water (b). LL: lower limit for soil water uptake.

The relationships between soil temperature and yield (Riley, 1957) in maize and between soil temperature and root growth in maize seedlings (Walker, 1969) have been known for half a century. However, the role of soil temperature during the growing season has been largely ignored as a determinant of maize productivity through resource capture and utilization. Geographical patterns in water uptake and/or depth of root presence in the soil profile, plausibly related to soil temperatures, could be constructed by comparing studies conducted across latitudes: 1.0m (Canada, Dwyer et al., 1996), 1.0m (Minnesota, USA; Fan et al. 2016), 1.2m (South Dakota, USA; Osborne et al., 2020), 1.3-1.5m (Iowa, USA; Ordóñez et al., 2018), 2.0m (Texas, USA; Tolk et al., 1998), and 2.1m (California, USA; Reyes et al., 2015). Because putative changes in root depth/occupancy have been shown to explain genotype (G) x environment (E) x management (M) interactions for yield in the US corn-belt (Hammer et al., 2009; Messina et al., 2011) it is logical to refine this hypothesis by stating: temperature-mediated increases in root occupancy and resource capture via water transport underpins GxExM interactions for maize yield. A corollary to this hypothesis is that these traits can contribute to yield improvement in temperate maize.

Root systems contain comprehensive mechanisms for tuning the water and nutrient capture relationships of crop plants through their development, function, morphology, architecture and interaction with the soil environment and microbiome (Reynolds et al., 2021; Karlova et al., 2021), and with the development of the leaf area (van Oosterom et al., 2016). Abiotic and biotic factors that limit the root system’s ability to grow and capture soil resources prevent a crop from reaching full productivity (Lynch and Wojciechowski, 2015; van der Bom et al., 2020). Direct selection for root system ideotypes and individual root traits is often hindered by phenotyping constraints combined with crop-level tradeoffs incurred when placed in the broader context of complex and varying agricultural production environments (van der Bom et al., 2020).

Public and private research and germplasm selection has focused on chilling stress resilience during seed germination and emergence to ensure stand establishment in cold, wet topsoil conditions and fluctuating springtime weather patterns (Menkir and Larter, 1987; Saab, 2013), however cold temperature isotherms persist within the soil profile throughout the growing season. These temperature isotherms are established and move through the profile based on daily and seasonal weather cycles and underlying soil texture, management and compositional properties (De Vries, 1963; Cruse et al., 1980; Kaspar and Bland, 1992). As roots grow through the soil profile they transect warmer to cooler temperature isotherms and push up against low temperature barriers that limit their growth, soil exploration and function (Stone et al., 1983). Root system exposure to sub-optimal temperatures have been found to reduce cell expansion (Pritchard et al., 1990) and cell division (Barlow and Adams, 1989b), reduce vessel diameters (Barlow and Adam, 1989a), and reduce root elongation (Pahlavanian and Silk, 1988) where growth ceases at temperatures below 10°C due to the disruption of sugar flow to the root (Crawford and Huxter, 1977). Lateral root initial abortion and changes embryonic root initiation angles and gravitropic responses of root meristems have also been documented. Additionally, impaired aquaporin function and decreases in root system respiration under low temperature reduce water and nutrient transport to the growing shoot and dynamically alter the carbohydrate sink localization within the root system (Onderdonk and Ketcheson, 1973; Sheppard and Miller, 1977; Atkin et al., 2000; Aroca et al., 2001; Hund et al., 2008; Nagel et al., 2009; Hund, 2010; Reimer et al., 2013; Lynch and Wojciechowski, 2015).

Together, low temperature isotherms are invisible factors that influence the extent to which root systems can explore the soil profile, uptake water and nutrients, and change the balance in resource allocation, all conducive to change the resource acquisition and utilization dynamics of the entire plant throughout the growing season. To gain insights on the effects of low temperature on root system growth and function, high-throughput phenotyping systems were developed, and experiments conducted to examine root systems’ responses to colder root zone temperatures. Genetic mapping was conducted and produced a nascent image of the genetic regulation underpinning root response to temperature in maize. Genotypic studies of water transport response to root temperature advanced our understanding of the degree of variation in the trait and can be implicated in the expression decreased in leaf water potential in the presence of available soil water. By integrating empirical and simulation results we discuss emerging opportunities to improve drought tolerance and maize productivity in the US corn-belt as mediated by changes in root systems form and function.

## Results

### Primary root growth rate decreased with decreasing temperature for hundreds of maize genotypes

Root growth response to low temperature was measured in a temperature-controlled growth platform capable of regulating shoot and root temperature independently (Fig. 2a,b). Roots were imaged (Fig. 2c) at regular intervals to estimate the primary root growth rate (PRG, mm d^-1^), and total root systems length and area growth rates (TRSG and TRSAG, mm d^-1^, and mm^2^ d^-1^) in response to changing temperature. On average, all root trait values decreased with decreasing temperature for maize hybrids and inbreds (Table 1). For temperatures below 18°C, the primary root growth for maize hybrids was not different from those of inbreds. A temperature dependent piecewise sigmoidal growth response curve was fit to the PRG for each genotype using a non-linear mixed model revealing almost no variation for a minimum base temperature of 8.54°C for root growth (TBmin, Fig 3a). A subsequent triangular cdf model was run with Tbmin fixed at 8.54°C and the maximum growth rate (RGRmax) ranged from 50.0 to 64.2 mm day^-1^ with an inflection point that varied between 18.6 and 19.6°C in maize hybrids with no significant correlation between traits (*r*^2^=0.003; Fig. 3a). Similar results hold for inbred lines with TBmin of 5.6°C and a range in rgrmax of 20.2 to 58.5 mm day^-1^ (Fig. 3b). The inflection point (TBmid) can be interpreted as a responsiveness to temperature where lower TBmid represents a lower responsiveness to decreasing root zone temperatures or an enhanced ability to grow under cooler soil temperatures.

**Figure 2.**
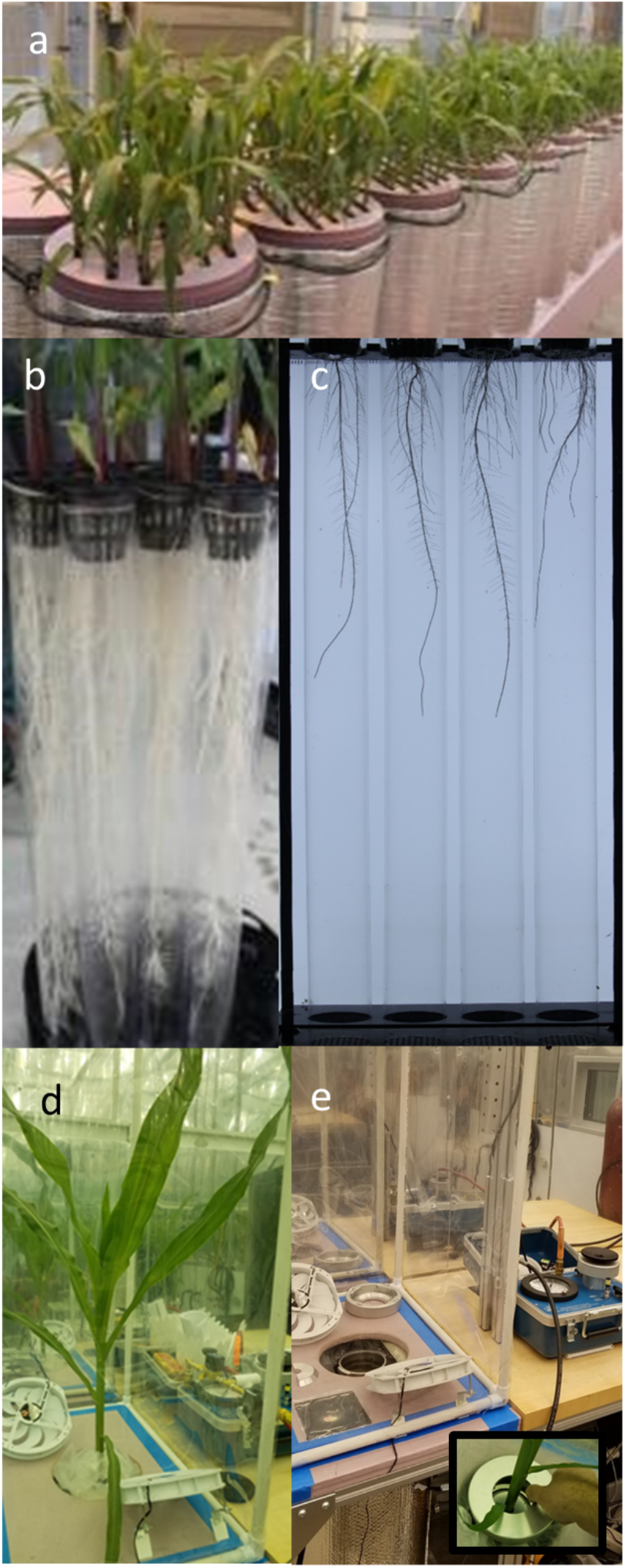
Temperature-controlled root phenomics platforms: 1) hydroponic root growth platform showing the temperature-controlled root growth platform module with growth tanks containing maize plants (a), a rack with individual growth tubes (b), and a raw image of root systems captured during root growth experiments (c), and 2) pressure chamber with root temperature control (d,e).

**Fig. 3.**
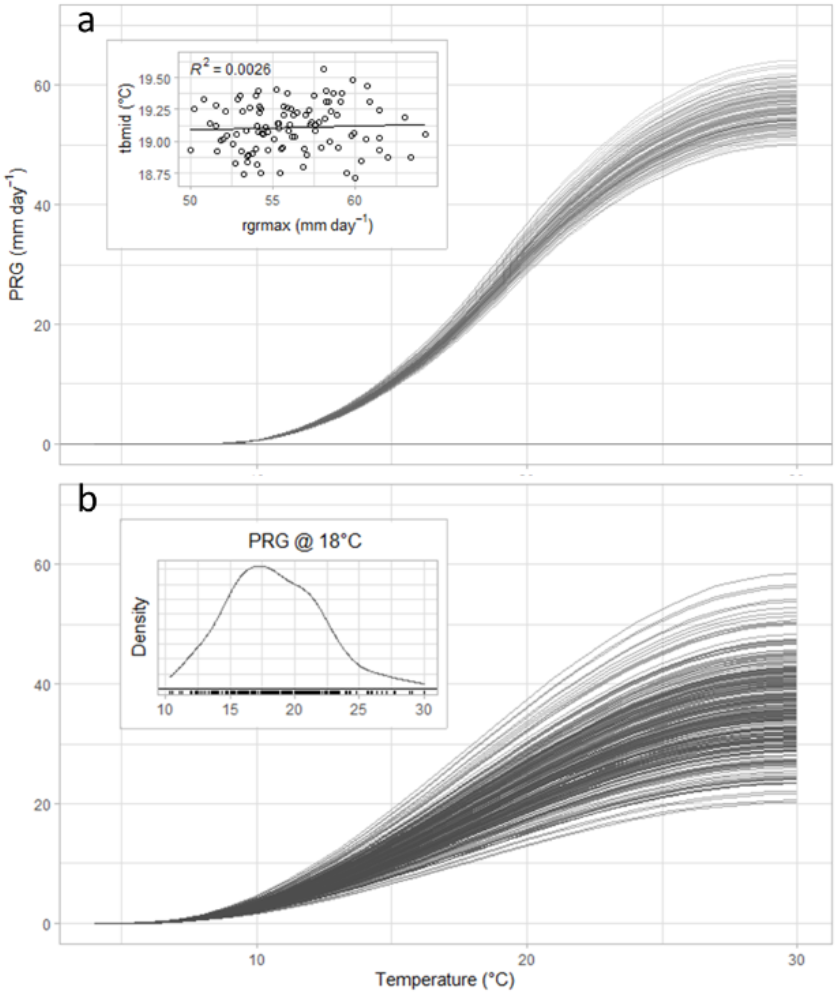
Growth response curves for maize genotypes across temperatures for primary root growth (PRG, mm day^-1^) with inset showing the relationship between inflection point (TBmid, °C) as a function of maximum rate of growth (RGRmax, mm day^-1^) across hybrids (a), and for inbreds (b) with inset showing the density function for PRG at 18°C.

**Table 1.**
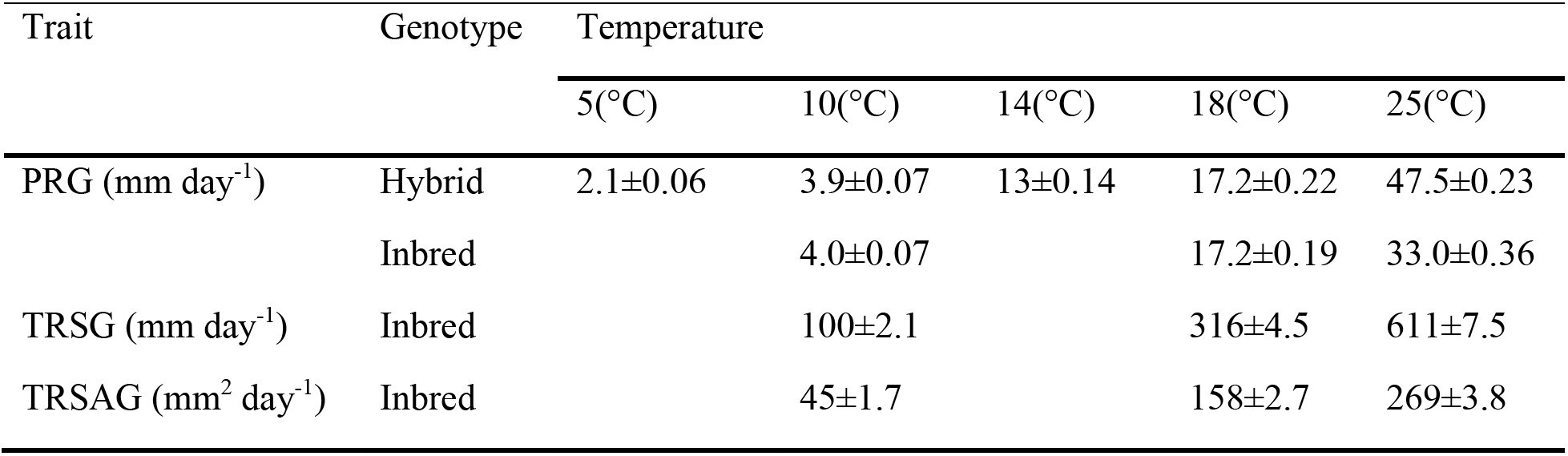
Average and standard errors for primary root growth rate (PRG), total root systems growth (TRSG), and total root systems area growth (TRSAG) by temperature for maize inbreds and hybrids.

### Root system conductance and root-shoot water flow decrease with temperature

A temperature-controlled root pressure chamber was developed to enable studying whole root system conductance response to temperature independently from shoot temperature (Fig. 2d,e). The system can accommodate plants with fully developed leaves undergoing C4 photosynthesis and enables the measurement of whole plant transpiration and carbon assimilation (Fig. 2d,e). For the hybrid P1498, light and transpiration response to photosynthetically active radiation were found to decrease with decreasing temperature (Fig. 4a,b). When root systems were acclimated to colder root zone temperature, maximum transpiration decreased from 6.0 to 3.8 mg H2O s^-1^ m^-2^, and photosynthesis decreased from 3.5 to 1.9 mg CO_2_ s^-1^ m^-2^ at 18 and 10°C, respectively. Consistent with this observation, the slope between transpiration and the balancing pressure (BP) decreased from 3.56 to 2.15 mg H_2_O s^-1^ m^-2^ MPa^-1^ (Fig. 4c) where a lower transpiration was observed for the same BP at low temperature. Root sap flow (RSF) also was significantly reduced from 7.6 to 3.2 mg H_2_O s^-1^ between treatments (Fig. 4c) in a similar to manner to the transpiration and BP responses indicating that lower leaf water potentials would be observed if the same whole root system water flow were to be maintained at a reduced root system temperature (Passioura, 1980).

**Figure 4.**
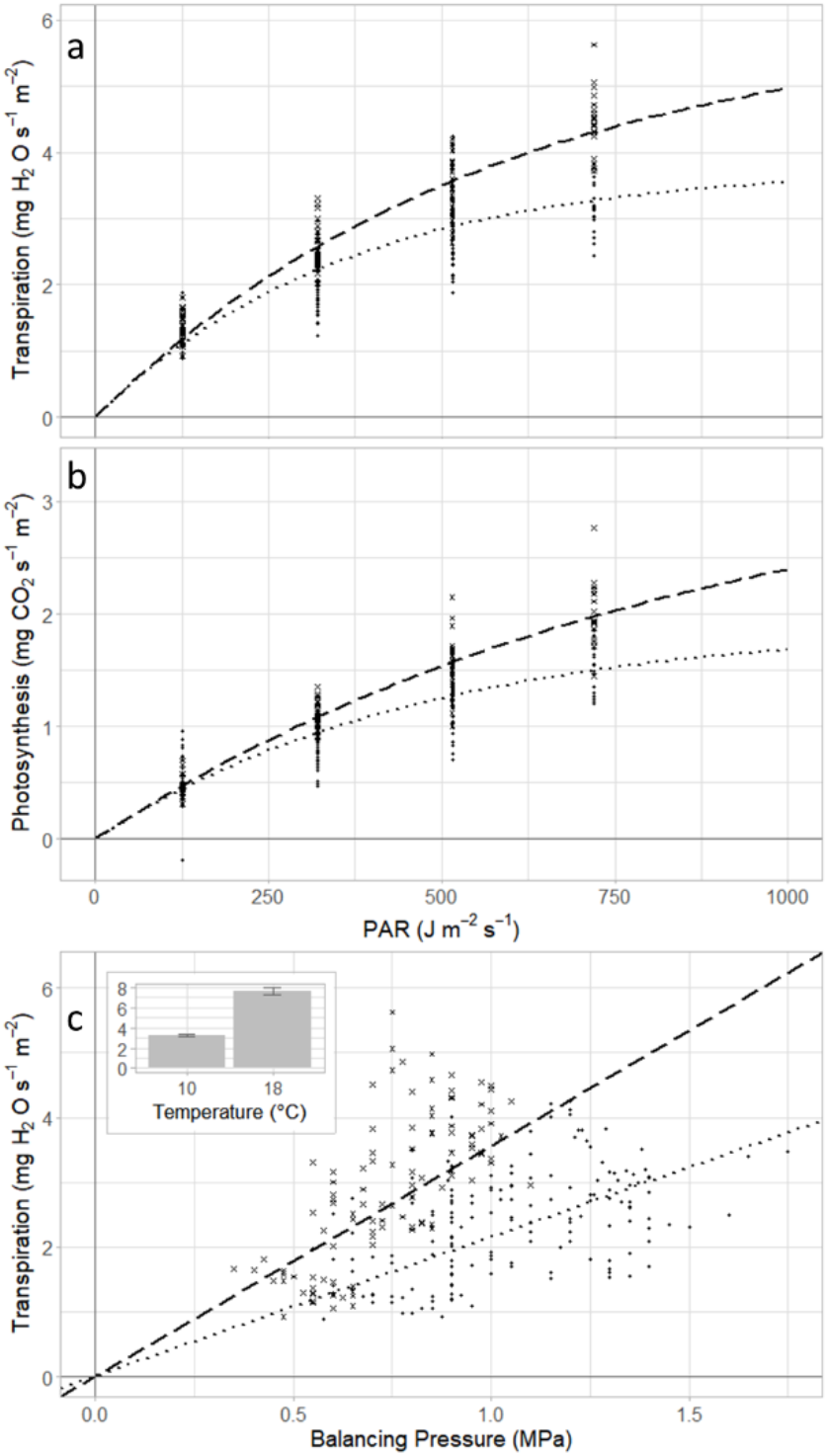
Whole plant transpiration to light (a), photosynthesis response to light (b), transpiration response to balancing pressure (c), and whole root system sap flow (inset (c), mg H_2_O s^-1^) is dependent on root temperatures set at 10°C (• and ⋯) and 18°C (× and ---).

### Root sap flow per unit leaf area varies between genotypes at constant low temperature

Root sap flow per unit plant leaf area (TPLA) was measured in 12 elite hybrids. At a whole root system temperature of 14°C RSF per unit TPLA significantly varied between hybrids from 31.6 to 36.5 mg H_2_O m^-2^ s^-1^ (Fig. 5). Because of the contrasting RSF per unit TPLA, the hybrids P0801, P1498, P1197, were tested at rooting temperatures 10, 15, 20, 25 and 30°C revealing that RSF was constant from temperatures 30 to 15°C but reduced to half of the maximum at 10°C. A linear plateau temperature response curve was proposed for subsequent simulation experiments and the temperature at the onset of RSF reduction when moving from higher to lower temperatures is defined as Tcond.

**Figure 5.**
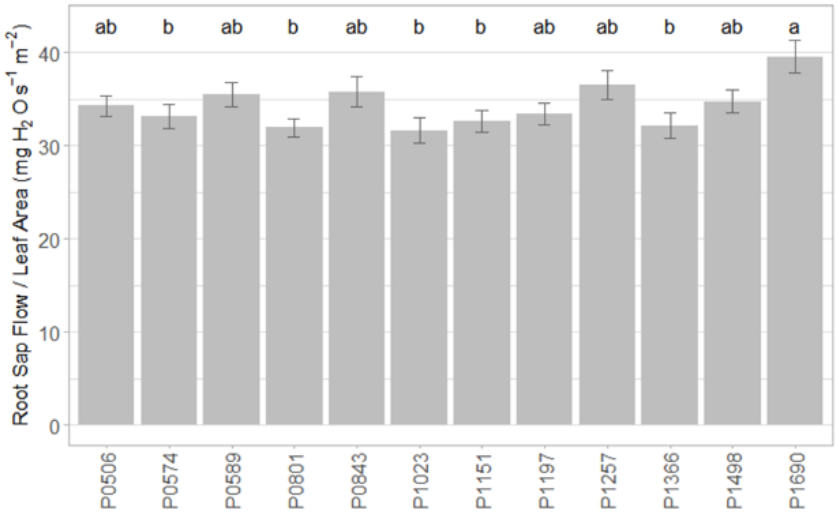
Genotypic variation for total root sap flow per leaf area (mg H_2_O s^-1^ m^-2^) for a set of elite maize hybrids treated at 14°C. Letter grouping indicates significant differences (p<0.05) using Tukey HSD.

### Genetic architecture of root growth under low temperature

To study the genetic architecture of root growth at low temperatures, genome wide association studies (GWAS) were conducted for inbred lines where PRG, TRSG and TRSAG were measured at three temperatures. Genetic markers were identified in significant association with each of the three traits, at one or more temperature conditions (Table 2). Specifically, three markers were detected for TRSG under all three temperature conditions (10, 18 and 25°C), and another marker was detected for TRSG at two temperature conditions (10 and 25°C). Otherwise, all markers were detected at only one temperature condition, and no markers were detected for multiple traits (Fig. 6). Certain of these markers were proximal to candidate genes with putative roles in cellular structure, root growth and stress tolerance. Candidates of particular interest include Zm00001d044760, which was proximal to a marker exhibiting significant association with TRSAG at 10°C. This gene product is annotated as TORTIFOLIA1-like protein 3 (https://www.ncbi.nlm.nih.gov/gene/?term=103639663) and has 72.9% similarity at the protein level with Os02g0739900/LOC_Os02g50640.1 in rice, which is annotated as a putative HEAT repeat family protein (Kawahara et al. 2013) and *tortifolia1-like protein 3* (https://www.ncbi.nlm.nih.gov/protein/XP_015623384.1). Another candidate of interest was Zm00001d030166/GRMZM2G321940, which was proximal to a marker exhibiting significant association with PRG at 18°C. The gene product is annotated as a glycosyltransferase-like KOBITO1 (Parvathaneni et al. 2020, Gramene).

**Figure 6.**
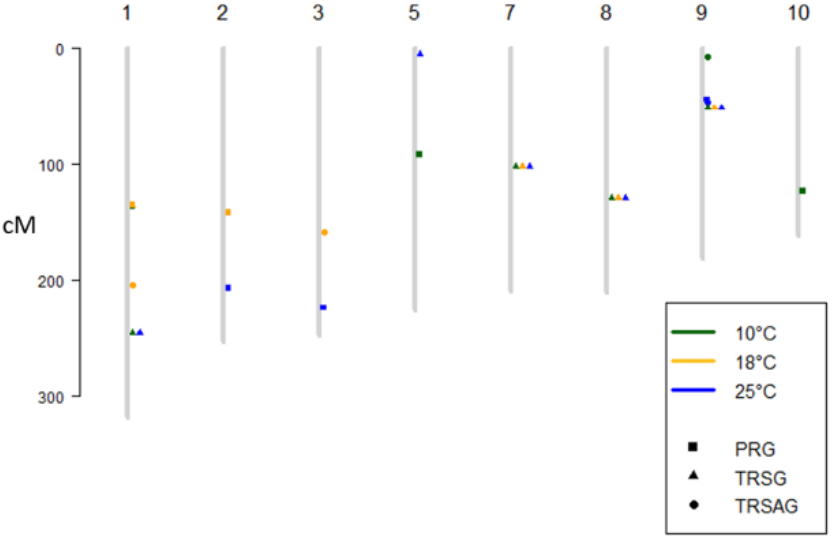
Genetic map with positions of markers associated with root growth traits dependent on temperature: Primary root growth (PRG), Total root system growth (TRSG), and total root system area growth (TRSAG).

**Table 2.** Market trait associations for primary root growth (PRG), total root system growth (TRSG), and total root system area growth (TRSAG) evaluated at three whole root system temperature, in the female (F) and male (M) side of heterotic group.

| Trait | T | Het.<br>Group | Marker ID: Name | Chr. | Pos. | $-\log_{10}(p)$ | Effect | $r^2$ |
| --- | --- | --- | --- | --- | --- | --- | --- | --- |
|  | °C |  |  |  |  |  | mm<sup>2</sup> d<sup>-1</sup> |  |
| TRSAG | 10 | M | C002PK7-001 | 9 | 5 | 4.1 | 36.0 | 0.18 |
| TRSAG | 18 | F | MZA10765-46 | 1 | 204.8 | 2.6 | -27.9 | 0.12 |
| TRSAG | 18 | M | C001MDD-001 | 3 | 159.4 | 2.1 | -61.4 | 0.10 |
| TRSAG | 25 | F | MZA10918-19 | 9 | 47.1 | 2.3 | -40.5 | 0.07 |
|  |  |  |  |  |  |  | mm d<sup>-1</sup> |  |
| TRSG | 10, 25 | F | C002Y3W-001 | 1 | 246.2 | 2.2 | -17.2 | 0.07 |
| TRSG | 10, 18,<br>25 | M | C0021E8-001 | 7 | 102.3 | 3.1 | 11- 18 | 0.14 |
| TRSG | 18 | F | MZA8982-3 | 1 | 136.5 | 2.1 | -19.2 | 0.06 |
| TRSG | 10, 18,<br>25 | F | C001X49-001 | 8 | 129.6 | 2.0 | -9.8 | 0.07 |
| TRSG | 10, 18,<br>25 | F | C00233H-001 | 9 | 52.1 | 2.3 | 18.8 | 0.10 |
| TRSG | 25 | F | MZA10373-7 | 5 | 5.5 | 2.3 | -12.4 | 0.08 |
|  |  |  |  |  |  |  | mm d<sup>-1</sup> |  |
| PRG | 10 | F | C001PGV-001 | 5 | 92.0 | 2.1 | -0.5 | 0.07 |
| PRG | 10 | F | C002GW1-001 | 10 | 124.1 | 2.4 | 1.1 | 0.06 |
| PRG | 10 | M | C001860-001 | 1 | 136.7 | 2.0 | -0.7 | 0.09 |
| PRG | 18 | F | MZA5336-25 | 1 | 135.9 | 3.4 | -1.8 | 0.14 |
| PRG | 18 | M | MZA4564-49 | 2 | 142.1 | 2.2 | -1.6 | 0.09 |
| PRG | 25 | F | MZA12969-14 | 3 | 224.4 | 2.3 | 1.5 | 0.07 |
| PRG | 25 | F | MZA11327-13 | 9 | 44.9 | 2.0 | -1.6 | 0.06 |
| PRG | 25 | M | MZA15127-20 | 2 | 207.1 | 2.3 | 2.1 | 0.10 |

### Impact of root systems response to temperature on yield: simulation assessment

To determine the manner and extent to which changes in root response to temperature affect yield across environments and in context of other physiological traits, an assessment was conducted using a simulation model (Cooper et al., 2104; Messina et al., 2015) where changes to root growth and function response to temperature were introduced (Fig. 3). The crop growth model used in this examination simulates both below and above ground physiological processes and their interaction with the environment (2012-2016 across the US corn-belt). Four cases were considered and compared to a baseline case for which parameters were set based on the experimental results shown above. Overall, a reduction of 1°C in TBmin, TBmid or Tcond increased average yields by 21, 18 and 1 g m^-2^, respectively. The simultaneous reduction in all three traits increased average yields in an additive manner by 40 g m^-2^. However, this yield benefit increased with decreasing environmental potential, mainly associated with water deficit (Fig. 7b). Yield improvements resulted from increased average root length density in depth (Fig. 7c) and consequently water uptake from the deeper layers within the soil profile (Fig. 7d). Geographical examinations by year indicate that yield improvements were neutral to positive in between 88 and 91 percent of simulations, depending on the trait/trait combination. In 2012, when water deficit was widespread throughout the US corn-belt, the combination of root traits demonstrated their potential contribution to attainable yield under water deficit conditions (Fig. 8). However, consistent yield reductions were also observed in some northwestern environments (Fig. 8). These yield reductions were likely due to unfavorable changes to the water capture dynamics, where rapid root growth in depth can increase water access and use during vegetative growth stages, thus lowering the available soil water during the critical reproductive stages (Fig. 8).

**Figure 7.**
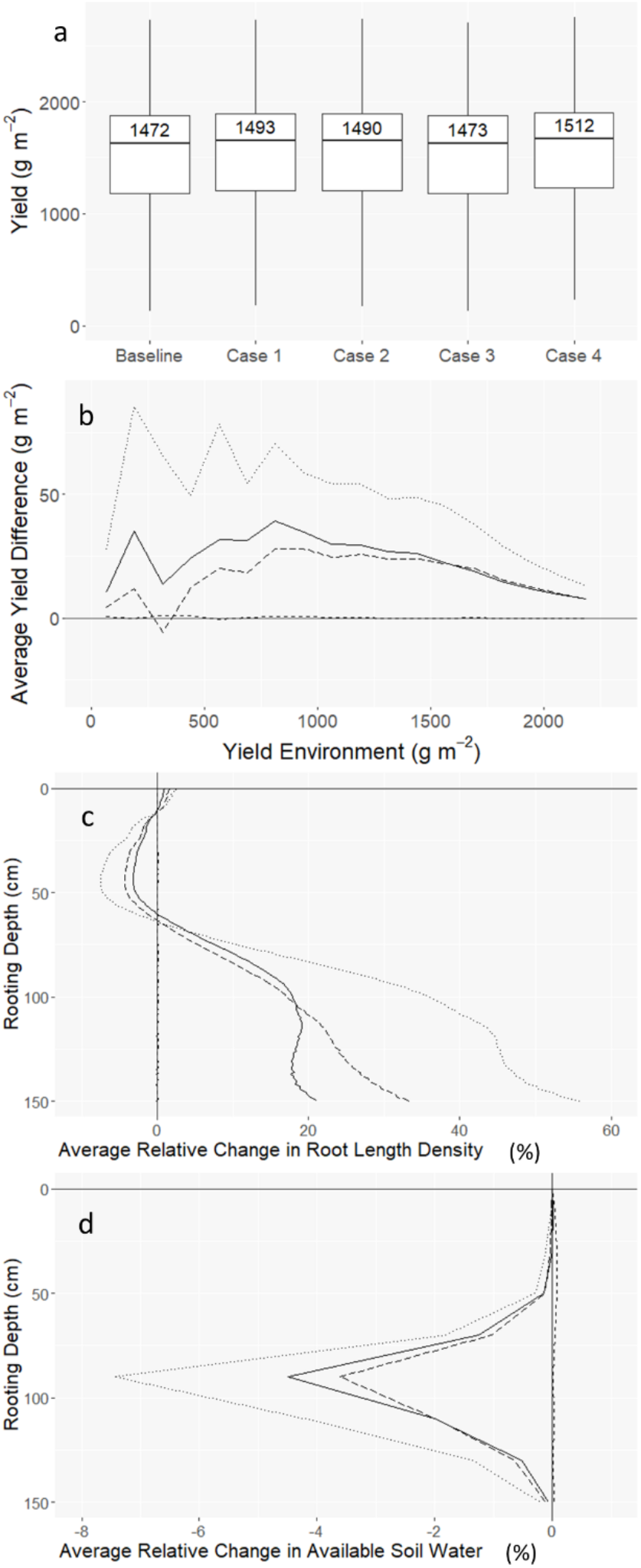
Simulated yields for 99 hybrids characterized using root phenomics (case 1: average TBmin lowered by 1°C, case 2: TBmid lowered by 1°C, case 3: Tcond shifted 1°C lower, case 4: decreases from cases 1-3 combined) for the period 2012-2016 in the US corn-belt vary slightly on average (a) but not on a productivity dependent manner (b, the four cases vs. baseline), which is related to the average relative change in root length density (c) and consequently on residual plant available soil water (d) with depth at flowering time (solid, long-dashed, short-dashed, and dotted lines represent Cases 1, 2, 3, and 4 respectively).

**Figure 8.**
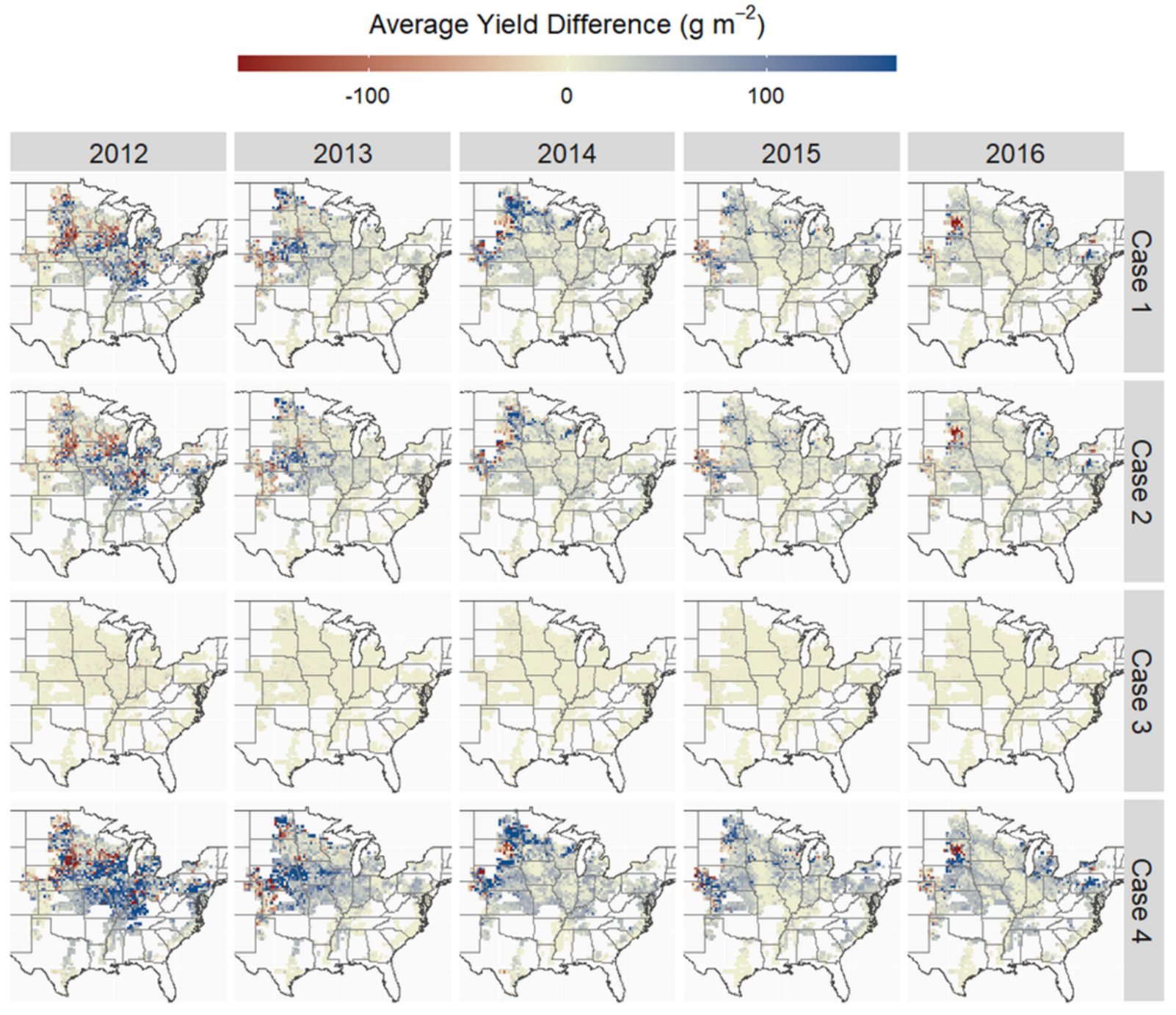
Average spatio-temporal yield differences between 99 baseline maize hybrids characterized for root response to temperature using root phenomic platforms with respect to hypothetical hybrids expressing root response to temperature phenomics (case 1: average TBmin lowered by 1°C, case 2: TBmid lowered by 1°C, case 3: Tcond shifted 1°C lower, case 4: decreases from cases 1-3 combined) for the period 2012-2016 in the US corn-belt.

To further examine how root responses to low temperature interact in the physiological background of the crop, an additional simulation experiment was conducted for a random sample of environments in the US corn-belt. The traits TBmin and TBmid were varied in a factorial manner and combined with each of 203 maize hybrids characterized for size of the ear leaf, leaf appearance rate, radiation use efficiency and its response to water deficit, conductance response to vapor pressure deficit, mass of the ear at first silk, total leaf number, and grain fill duration (Messina et al., 2020). Figure 9 shows the relative contribution of each physiological trait to the simulated yield, conditional upon the environmental potential. The contributions of TBmin and TBmid to yield was highest for yield environments ranging from 600 and 1400 g m^-2^, which is also the range of environments where yield determination results from contributions from many traits and their interactions.

**Figure 9.**
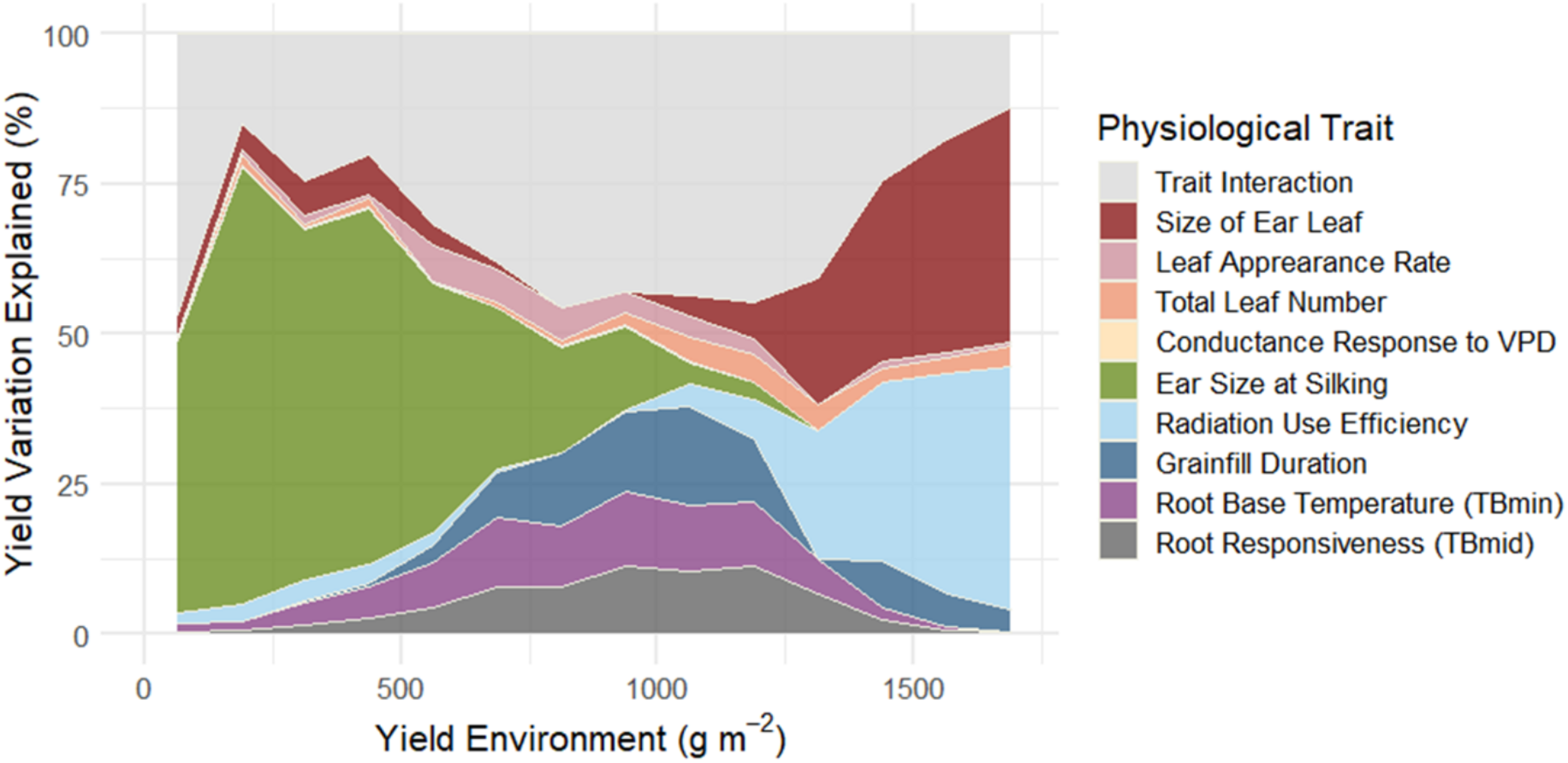
Average relative contributions of reproductive traits, root and shoot to yield across a productivity gradient of environments for 203 elite maize hybrids simulated with minimum base temperature for root growth (TBmin) ranging from 8.54 to 4.54°C and root growth response to temperature inflection point (TBmid) between 19.6 to 15.6°C on 200 sites located within the US corn-belt between 2012-2016.

## Discussion

Although the effects of soil temperature on root growth of maize seedlings (Walker, 1969) and yield (Riley, 1957) have been known for more than half a century, there has not previously been evidence that root growth and function response to temperature underpin genotype x environment interactions, nor have studies indicated how to harness this knowledge to inform crop improvement in maize by increasing water capture. Through a combination of physiological, genetic and simulation studies, we provide a nascent view of the genetic regulation of whole root systems response to temperature in maize beyond the seedling stage. Root temperature was found to limit transpiration and photosynthesis response to photosynthetically active radiation, and genotypic variation was identified for the low temperature growth and conductance traits in modern germplasm. We also demonstrate that root systems occupancy is far more important than conductance in the determination of yield across a wide range of yield environments. The contributions of the low temperature growth responses traits were greatest in yield environments characterized by yields between 600 and 1400 g m^-2^. Strikingly, these yield environments represent where most of the US production acres lie. This knowledge helped advanced our understanding of root biology and yield determinants in maize and created the opportunity to inform crop improvement through mathematical prediction (Messina et al., 2018; Cooper et al., 2020; Messina et al., 2022c). Selecting germplasm for improved capacity to access water will be necessary to continuing harnessing improvements in radiation use efficiency (Messina et al., 2022b). In an increasingly warmer, drier, and volatile climate, our results can open an opportunity to sustain crop improvement to water deficit (Cooper et al., 2014; Messina et al., 2022a) and improve the adaptation of maize and other summer crops to climate change (Cooper et al., 2021; Cooper and Messina, 2023).

### Whole plant phenotyping

This research was enabled by constructing unique phenotyping systems to study temperature controls for the root and shoot system separately for plants at development stages past the seedling stage. This platform improves upon prior integrated systems (Clark et al., 2011). While in the present study we focused on the elongation for the primary root (e.g. as in Pahlavanian and Silk, 1988), the total root system, and the area occupied by roots, other traits could be measured to continue progressing our understanding of how low temperature ultimately regulates growth, transpiration and yield. Image analysis algorithms could be expanded to study genetic variation in root branching and abortion of lateral root initials (Barlow and Adam, 1989a), root initiation angle, and gravitropic responses (Onderdonk and Ketcheson, 1973; Sheppard and Miller, 1977; Atkin et al., 2000; Aroca et al., 2001; Hund et al., 2008; Nagel et al., 2009; Hund, 2010; Clark et al., 2011; Reimer et al., 2013; Lynch and Wojciechowski, 2015). The balancing pressure systems in combination with reducing sugar analyses could provide insights on genetic variation for sugar flow to the root (Crawford and Huxter, 1977) and how soil temperature can alter the dynamic allocation of carbon in the plant and within the soil profile. In addition, experiments that combine temperature, water deficit and genotype treatments can lead to further understanding of how hormonal signals mediate root/shoot allocation in response to soil temperature and water potential. Overall, the platform described in this study can be instrumental to translate root science into breeding goal by at least partially removing a critical bottleneck in root and crop physiology (Reynolds et al., 2021).

### Rooting depth

Many empirical and simulation studies have suggested that deep rooting to access water supplies lower in the soil profile and improve yield under drought stress (e.g., Hammer et al., 2009; Messina et al., 2011; Lynch, 2013; Lynch et al., 2014; Reynolds et al., 2021). Lower branching and metabolic costs (Lynch et al., 2014; Zhan et al., 2015) and adaptive root response to soil water (Orosa-Puente et al., 2018) have been implicated in the development of deep root systems. Recent studies further our understanding to link the multiseriate cortical sclerenchyma phenotype to root penetration in compacted soils (Schneider et al., 2021). Our results offer an additional explanation for increased rooting rate, mass production and plausible yield. Considering that low temperatures increase abortion of lateral root initials (Barlow and Adam, 1989a) it could be interesting to test whether the lower branching phenotype is associated with deep rooting or if the abortion of lateral initials is an adaptive strategy to expand soil exploration under increasing cooler isotherms.

### Genetic architecture of root response to temperature

Independent of root system vigor, root system responsiveness to low temperature was shown to vary within hybrids, and this variation appears to influence overall rooting depth and length density which were found in a simulation study herein to result in yield improvement across large regions of the US corn-belt. The GWAS conducted on inbred lines revealed that independent genetic loci can act at different temperatures suggesting that some portion of these changes can be genetically selected for and optimized within regions of the growth response curve. Root sap flow and conductance show that there is a genotypic basis to root conductance responses to decreasing temperature. We propose it should be feasible to leverage natural variation in all traits identified in this study to hasten genetic gain for yield using prediction approaches (Diepenbrock et al., 2022; Messina et al., 2022c; Cooper and Messina, 2023).

Gene editing of candidate genes can contribute to speed genetic gain for yield. Here, we identified such candidates. The protein encoded by Zm00001d044760, proximal to a marker detected for TRSAG at 10°C, exhibited high similarity with Os02g0739900/LOC_Os02g50640.1 in rice. This protein in rice is annotated as a putative HEAT repeat family protein (Kawahara et al. 2013), and has been noted in the context of relatedness to the plant-specific microtubule-associated protein (MAP) family containing *tortifolia1/spiral2,* though with only 27% protein identity to *tortifolia1/spiral2* in Arabidopsis (Guo et al. 2009). *tortifolia1/spiral2* in Arabidopsis is indeed a plant-specific MAP containing HEAT-repeat motifs, and recessive mutation results in right-handed helical growth and relatively mild (compared to another helical growth mutant, *spiral1)* defects in growth anisotropy, including in roots (Buschmann et al. 2004, Shoji et al. 2004, Furutani et al. 2000). While *spiral1* was found to have a more pronounced mutant phenotype at low temperature that was nearly completely suppressed at higher temperature, *tortifolia1/spiral2* was previously found not to exhibit that temperature dependency in Arabidopsis (Furutani et al. 2000). However, the marker proximal to Zm00001d044760 having been detected at 10°C in the present study suggests that further screening of variants of this gene at low temperatures in maize may be informative, including for purposes of breeding for increased TRSAG at low temperatures. The protein encoded by Zm00001d030166, proximal to a marker detected for PRG at 18°C, is annotated as glycosyltransferase-like KOBITO1 (Parvathaneni et al. 2020, Gramene). In Arabidopsis, *kobito1* (characterized in Pagant et al. 2002) is allelic to *abscisic acid-insensitive 8 (abi8;* Brocard-Gifford et al. 2004) and *elongation defective1 (eld1;* Cheng et al. 2000, Lertpiriyapong and Sung 2003). These mutants have been characterized in Arabidopsis with observed defects in cell elongation (in multiple organs, including roots) and PRG (Cheng et al. 2000), as well as observed cellulose deficiency (Pagant et al. 2002). A marker proximal to Zm00001d030166 having been detected in this study for PRG suggests that this candidate gene could merit further investigation in maize.

### Root traits determination of crop adaptation

In plant breeding, adaptation is often considered for one trait dimension at a time – examples include root depth, water flow, root penetration, partitioning, and other scientific bottlenecks to yield improvement (Reynolds et al., 2021). In the present study, we show that the impacts of root trait variation are dependent on both the environment and the physiological-genetic context that determine the state of traits of the genotype. In extremes of production environments, rooting was not found to have a major impact on simulated yield for the conditions of the US corn belt. In extreme drought, the lack of water available in the soil dictates a null effect of the increase root exploration. In water sufficient environmental conditions, increase soil exploration is not necessary to capture water to satisfy the crop water demand. Instead, it is in the most typical production environments that root trait variation was found to have the most consistent yield benefit. These environments encompass a mixture of intermittent water deficits, punctuated deficits at flowering time, and extended periods during grain fill (Löffler et al., 2005; Messina et al., 2015). Testing the hypothesis proposed by Hammer et al. (2009), it was unexpected to find that long-term selection for yield did not contribute to shift rooting depth (Reyes et al., 2015; Messina et al., 2021), despite simulation studies suggesting the contrary (Hammer et al. 2009; Messina et al., 2011). Consistent with the contributions of root systems response to temperature on yield being dependent on other traits, one hypothesis could be that long-term selection improved reproductive resilience and that up to now the physiological background of modern US maize has not been conducive to the expression of the benefits of a deeper root system on yield. Chapman et al. (2003) reported the potential for similar conditional and sequential contributions of traits for long-term yield gain of sorghum in Australian dryland environments. Diepenbrock et al. (2022) using a combination of empirical yield trials and simulation propose that an important component of genetic variation for yield in modern maize hybrids could be explained by rooting traits. Future studies should include larger populations to increase the power to further detect markers associated with root traits in elite maize.

## Conclusion

There is genetic variation for root system response to temperature and genotypic variation in conductance response to temperature in temperate maize. The GWAS analysis detected marker-trait associations for the herein examined root traits. While soil temperature affects water conductance and transpiration, it is not immediate an effect of yield based on simulation assessment. In contrast, root growth in depth and occupancy offer a nascent opportunity to improve yield and drought tolerance in maize by harnessing the knowledge presented in this research.

## Materials and Methods

### Root growth response to temperature phenotyping

Root growth experiments were conducted on a maize (*Zea mays* L.) inbred diversity panel and a modern maize hybrid panel consisting of 249 lines and 99 hybrids, respectively. Plants were evaluated in a temperature-controlled, hydroponic root growth and imaging platform within the controlled environment greenhouses at Corteva Agriscience in Johnston, IA. The growth platform consisted of individual modules, each containing an insulated 760 L supply tank, 40 insulated 57 L growth tanks, a centrifugal water pump, a water heater and chiller, a PLC control unit, and component plumbing, wiring, and temperature and flow sensors. The growth tanks were arranged into 4 tank sets of 10 growth tanks each and a modified Magnavaca’s nutrient solution (Magnavaca et al., 1987) was supplied to each tank set on a regular cycle via pump-assisted ebb and flow. The growth modules and tanks contained an integrated misting system that was connected to the supply tank for supplemental temperature control in-between ebb and flow cycles between tank sets. Inside each growth tank was a rack with 22 transparent plastic growth tubes 56 cm height x 3 cm diameter open-bottom growth tubes with a netpot at top filled with rockwool for growing the plants individually and facilitate temporal imaging of their roots. During the growth experiments, pre-germinated seedlings with primary roots between 3 and 10 cm long were transplanted into the growth tubes and pre-grown with a root zone temperature of 25°C for 5 days for inbred experiments and 4 days for hybrid experiments. For inbred experiments, the pre-grown plants were imaged immediately prior to being moved into treatment modules that were held at 10, 18 or 25°C, then imaged again after 3 days. For hybrid experiments, the pre-grown plants were subjected to a 3-phase temperature course of either 25:18:10 or 25:14:5°C. Images of the root systems were captured at the beginning and/or end of each phase of the treatment course where phase one, two and three lasted 2, 2 and 3 days, respectively. The root system images were captured on a custom imaging system and analyzed with custom MVTec HALCON HDevelop (Eckstein and Steger, 1999) and Fiji (Schindelin et al., 2012) programs and plugins. From the captured root systems images, total root system area (mm^2^), total root system length (mm) and primary root length (mm) were measured and the sequential measurements for each plant were used to calculate the total root system area growth (TRSAG in mm^2^ day^-1^), total root system growth (TRSG in mm day^-1^) and primary root growth (PRG in mm day^-1^) rates.

### Root growth response to temperature statistical analyses

Growth rate BLUPs within each temperature (10, 18 and 25°C) for each inbred were estimated using linear mixed models

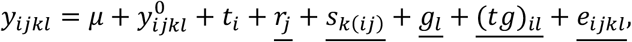

where *y_ijkl_* is root growth rate for inbred *l* from replication *j* and rack set *k* at temperature *i, μ* is the overall mean, 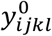 is the root length or area at the beginning of treatment, serving as a covariate, *t_i_* is the main effect of temperature *i, g_l_* is the main effect of inbred *l*, (*tg*)_*il*_ is the interaction effect of temperature *i* and inbred *l, r_j_* is the effect of replication *j, s*_*k*(*i,j*)_ is the effect of racket set *k* from temperature *i* and replication *j* combination, and finally *e_ijkl_* is the residual effect. All underlined terms are assumed to be normally distributed random terms with mean 0. The models were fitted using ASReml (Gilmour et al., 2009).

Individual growth coefficients (TBmin and RGRmax) for each inbred and hybrid were estimated from nonlinear mixed models with the underlying nonlinear function a piecewise sigmoidal growth curve

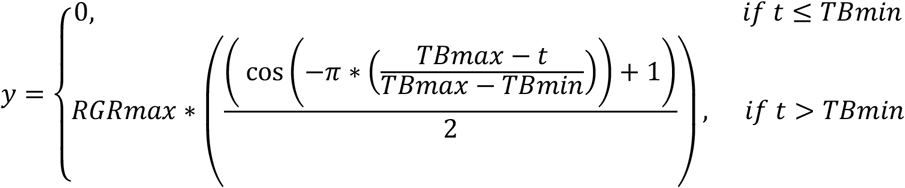

The temperature of maximum growth (TBmax) was fixed to 30°C based on finding from Kaspar and Bland (1992) while the temperature of minimum growth (TBmin) and maximum root growth rate (RGRmax) depended on initial root length and genotype. Taking TBmin as an example, the model is,

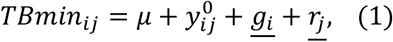

where 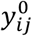 is the initial root length at the beginning of treatment for genotype *i*, replication *j* and *g_i_* is the random effect of genotype *i* and *r_j_* is the random effect of replication *j* for the hybrid experiment. Models were fitted using the nlme package within R (R Development Core Team, 2010; Pinheiro and Bates, 2020).

Using the estimated TBmin coefficient, the maximum root growth rate (RGRmax) and inflection point (TBmid) for the hybrids were then estimated from a nonlinear mixed model with the underlying nonlinear function a piecewise triangular cumulative distribution function curve,

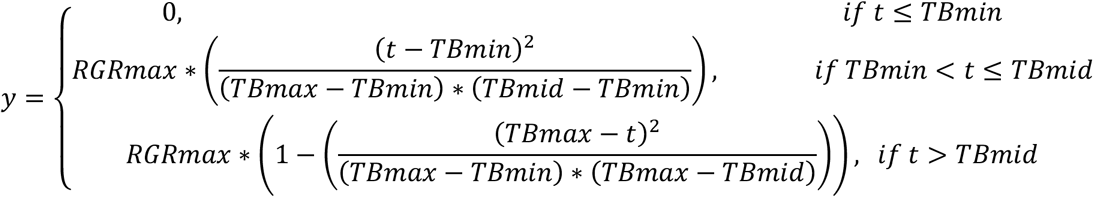

where the RGRmax and TBmid depended on initial root length and genotype and were modelled using the same model as model (1).

### Water flow response to whole root system temperature

Root function and conductance experiments were performed on maize hybrids using a temperature-controlled root pressure chamber system. The root pressure chamber system consisted of a Model 600-EXP Super Pressure Chamber from PMS Instrument Corporation with additional temperature control and a reengineered sealing orifice to accommodate whole plant stalks up to 22 mm diameter. The root pressure chamber was integrated onto a mobile cart with a transparent film shoot enclosure made with polyethylene terephthalate film with internal and external fans, wiring and pumps, a CR1000 datalogger from Campbell Scientific, and LI-840 CO_2_/H_2_O gas analyzer from LI-COR Biosciences.

All plants were grown in the greenhouse in PVC tubes containing general purpose potting substrate composed of peat moss, vermiculite, starter fertilizer and Osmocote. Prior to testing when the plants had reached a V4-V6 growth stage, the tubes were temporarily sealed at the bottom with plastics bags and moved into a walk-in growth chamber containing temperature-controlled water baths where the plants could acclimate to root temperatures of 10, 14 or 18°C for 2 nights prior to testing. All plants were grown under well-watered conditions throughout the acclimation period with growth chamber settings of 29°C/23°C (day/night) with 450 J m^-2^ s^-1^ light level and no added humidity.

For light response studies, a single commercial hybrid (P1498) was selected for testing and was grown in 60 cm height x 8 cm diameter mesh-bottom tubes. The plants tested at separate light levels of 125, 320, 515 or 720 J m^-2^ s^-1^ with root temperatures of 10 or 18°C with between 15 and 26 replicates per light level and treatment. Once the plants root system was sealed into the temperature-controlled pressure chamber and the shoot was isolated, an automated testing program allowed for whole plant transpiration and photosynthesis to be recorded over a 43-minute period prior to pressurizing the roots to observe the balancing pressure (Passioura, 1980) where the balancing pressure was measured as the lowest pressure within the root pressure chamber when a stable-size, non-dripping droplet of guttation was observed and held on the tip of the youngest fully-expanded leaf. The program consisted of a 20-minute acclimation period where an external refreshing fan moved outside air into the enclosed shoot chamber preceding 6, 2-minute, observation periods where the refreshing fan was stopped, covered, and the change in H_2_O and CO_2_ was recorded. Each 2-minute observation period was separated by a 1-minute refresh period to allow fresh air in.

After recording the balancing pressure, and for the root system conductance only experiments, the shoots of the plants were cut off at 10 cm above the base of the stalk. The cross-sectional area of the stalk at the cut was measured with digital calipers and total leaf area of the plants was measured with a LI-COR LI-3100C area meter. Xylem sap exuding from stalks (termed root sap) was collected for 5 minutes while keeping the root systems under 0.5 MPa of pressure. Sap was collected by placing a pre-weighed, conical falcon tube with tissue paper to absorb and contain the root sap. For the root conductance only experiments, the plants were grown in 30 cm height x 4 cm diameter, open-drained tubes and 12 commercial hybrids (P0506, P0574, P0589, P0801, P0843, P1023, P1151, P1197, P1257, P1366, P1498, P1690) were selected for testing at a root temperature of 14°C with between 21 and 27 replicates per hybrid. Because absolute phenotypic differences among genotypes, and thus the genotypic signal/noise ratio, decrease with decreasing temperature, a preliminary study was conducted with a subset of 4 of the 12 hybrids. These were tested at 10, 12 and 14°C to determine the temperature that would enable an effective separation of hybrids (unpublished data).

### Genome wide association studies (GWAS)

Genome wide association studies were conducted on TRSAG, TRSG, and PRG for each temperature treatment (10, 18 and 25°C) using 241 of the inbred lines that were genotyped with 8642 genetic markers. A 100-iteration permutation test with 5th percentile -log10(p) selection threshold was used to determine significant makers. Due to extensive population structure separate GWAS analyses were conducted within heterotic groups, with 102 lines from the male side of the pedigree and 122 inbred lines from the female side. Candidate genes were identified within the search space (±1 marker of the marker showing signal) in addition to MaizeGDB and TAIR searches (Swarbreck et al., 2007; Portwood et al., 2018).

### Simulation assessment of root response to temperature traits

For simulations studies, a stochastic root system architecture model was integrated with a crop growth model that was previously described (Cooper et al., 2014; Messina et al., 2015). The root system architecture model was written in Java (Arnold et al., 2005) and was designed building from root growth and development modeling principles (Pellerin, 1993; Pagès et al., 2000; Lobet et al., 2015) with specific functionality to allow the root systems to respond daily to localized soil conditions and phenological outputs from the crop growth model. Simulations studies of temperature response across the US corn-belt were conducted between 2012 and 2016 across the US corn belt (Löffler et al., 2005; Messina et al., 2015) under non-irrigated conditions to sample varying weather patterns within production geographies of the US and four case studies were designed to evaluate and compare the measured hybrid population from the growth experiments (Baseline) to a set of hypothetical populations with improved low temperature root growth and/or function responses, totaling approximately 16.5M simulations. The low temperature response improvements were as follows, for Case 1 the minimum base temperature of growth (TBmin) was lowered by 1°C; for Case 2 the responsiveness (TBmid) was lowered by 1°C; for Case 3 the linear plateau conductance response curve and the temperature at the onset of the conductance reduction (Tcond) was shifted 1°C lower; and for Case 4 the growth and conductance improvements from Cases 1, 2 and 3 were combined. To further investigate relative importance and interaction of targeting root traits on currently known shoot traits, a sample of 203 single cross commercial and precommercial mid-maize hybrids with previously measured crop growth model traits were selected and simulated with an RGRmax of 57.1 mm day^-1^ and TBmin and TBmid coefficients ranging from 4.54 to 8.54°C and 15.6 to 19.6°C by every 1°C. Simulations were run over 200 randomly selected locations in the US corn-belt between 2012 and 2016 to equally sample a variety of yield level environments ranging from 0 to 1800 g m^-2^. ANOVA was then performed on the predicted yield results and the relative contributions of TBmin, TBmid and the other shoot traits were plotted across yield level environments. All together 5075 individual hybrids were modeled with 1 replicate model run per hybrid, totaling 5.1M simulations. Weather data used to run the model were from NOAA, soils data were from USGS, and agronomic management practices were defined based on prior publications (Messina et al., 2015; Cooper et al. 2020).

## Data

At the sole discretion of Corteva Agriscience the data could be made available upon request.

## Acknowledgements

We thank Steven Becker and Ross Megargel for their help with engineering and fabrication of the phenotyping platforms, Karen Thompson, Katie Strand, Joshua Snoberger and Logan Anderson for their assistance with experimentation, Yinan Fang for his help with statistical modeling, and Jose Rotundo for his discussions and feedback when preparing the manuscript. William S. Niebur and John Arbuckle for their support over two decades of research at Pioneer Hi-bred and Corteva Agriscience. C.M. receives support from the IoT4Ag Engineering Research Center funded by the National Science Foundation (NSF) under NSF Cooperative Agreement Number EEC-1941529. Any opinions, findings and conclusions, or recommendations expressed in this material are those of the author(s), and do not necessarily reflect those of the NSF.”

